# LICHEN: Light-chain Immunoglobulin sequence generation Conditioned on the Heavy chain and Experimental Needs

**DOI:** 10.1101/2025.08.06.668938

**Authors:** Henriette L. Capel, Isaac Ellmen, Chris J. Murray, Giulia Mignone, Megan Black, Brendan Clarke, Conor Breen, Sean Tierney, Patrick Dougan, Richard J. Buick, Alexander Greenshields-Watson, Charlotte M. Deane

## Abstract

In developing therapeutic antibodies, the heavy chain is often prioritised due to its higher variability and its central role in antigen binding. An appropriate pairing of the light sequence is however important for antibody function. Here we present LICHEN, a heavy chain conditioned light sequence generation tool that enables collaborative light sequence design by leveraging computational capabilities alongside experimental expertise. LICHEN generates light sequences which are valid (antibody-like), diverse in sequence and structure, and conditioned on a specific heavy chain. LICHEN can also condition on germline and CDRs and automatically filter generated sequences for required properties. This allows LICHEN to be used across multiple antibody development use cases. We carry out experimental validation of the method conditioning only on the heavy sequence and on the heavy sequence and binding information. Our *in vitro* results show that sequences created by LICHEN have effective expression yields and can retain antigen-binding.

## Introduction

Antibodies are essential proteins of the adaptive immune system. The specific binding of the antigen with high affinity, coupled with the potential to further mutate in response to a target makes antibodies an excellent potential therapeutic class(1). A monoclonal antibody consists of two identical heavy chains paired with two identical smaller light chains(2). The complementarity determining regions (CDRs), located at the tips of the variable region (Fv), contain most variability and mediate antigen binding. The variable heavy (VH) and light (VL) chain contain three CDRs each, with CDRH3 being the most variable.

Diversity of the CDRH3 derives from the combinatorial and junctional diversity of the three genes (V, D, and J) encoding the full VH sequence. The VL sequence is encoded by only the V and J-gene. During B-cell development, the VH gene locus is first rearranged, after which the VL chain is rearranged to produce a suitable light chain that pairs with the heavy chain(3). Upon encountering a specific antigen, these germline encoded “naive” VH-VL sequences undergo an affinity maturation process which creates sequences that bind strongly to that antigen(3).

Modern antibody discovery for therapeutic purposes often starts with library-based discovery(4). Generative language models, such as IgLM(5) and p-IgGen(6), allow the generation of large synthetic antibody libraries. IgLM and p-IgGen are both decoder-only models, originally designed for text generation tasks(7). IgLM is trained on single chains, species and chain type. p-IgGen is trained on paired data after pretraining on unpaired data and can therefore generate paired VH and VL sequences. Developability issues for antibodies generated by language models have been reported (8) and these models are unaware of the antigen. Thus, generic libraries designed by these methods require optimisation for affinity to the target antigen, and developability during downstream antibody design pipelines.

In the development of antibody therapeutics there has been a focus on the heavy sequence due to its greater variability(9) and its importance in binding(10). This is exemplified in the availability of unpaired sequence data. For example, there are six times as many VH sequences as VL sequences in OAS^1^(11). However, it is known that the proper pairing of the light chain to a heavy sequence is essential for the functionality of an antibody(12–14).

Constructing the appropriate light sequence for a given heavy chain could be done using generative models, especially when combined with prior attained knowledge on key regions for antigen binding or germline usage. This maximises the potential of the computational tools to guide, and experimental knowledge to inform the antibody engineering workflow. Here, we present LICHEN, a light sequence generation tool tailored to a given heavy sequence and (if desired and/or available) to experimental prior knowledge. LICHEN is a sequence-to-sequence model trained on natural paired human Fv sequences. We demonstrate that generated sequences are valid human light sequences, which are diverse and a fit for the heavy sequence, simulating patterns observed in nature. In addition to the heavy sequence, LICHEN flexibly supports conditioning on CDR sequences and germline information, and offers applications when only heavy sequence information is available (e.g. from unpaired heavy sequencing), as well as when additional binding information is available (e.g. sequences derived from animal immunisation). Germline guidance improves applicability to experimental settings (e.g. when specific V-gene usage is preferred). This is further enhanced by automatic sequence filtering options. *In vitro* we show that light sequences generated by LICHEN express well and maintain antigen-binding when CDR information is provided. LICHEN is available open source as both a python package (https://github.com/oxpig/LICHEN) and web application (https://opig.stats.ox.ac.uk/webapps/lichen).

## Methods

### Dataset generation

Paired human VH-VL sequences were curated from the publicly available database OAS(11) in November 2023. Sequences missing the conserved cysteines in IMGT positions 23 and 104 were removed, as well as sequences with unknown residues and duplicate pairs. Deletions in framework (FR) residues, observed in around 8% of the antibody pairs, were restored with AbLang(15). The remaining data was randomly split into 80% training, 10% validation and 10% testing. Only exact duplicate heavy sequences were excluded from the test set. The filtered and cleaned data for training and testing have been deposited on Zenodo (https://doi.org/10.5281/zenodo.15917096). Training was performed on 1,452,229 paired sequences.

The performance of LICHEN was tested on a random subset of the test set, the “standard_ds”. We selected 500 test set samples and generated 20 light sequences for each heavy sequence. When not specified otherwise, the standard_ds is used for evaluation. For structural evaluation we selected five heavy sequences from five different V-gene families which all naturally paired with more than 20 light sequences, the “structural_ds”. We sampled and generated 20 light sequences for these heavy sequences. The conditioning ability of LICHEN was evaluated on the “conditioning_ds”. We selected 1000 test set samples each with between five and fifteen mutations away from the germline in both chains. Germline reverted light sequences were constructed by mutating the identified mutations back to the closest germline Vgene and J-gene (as determined by ANARCI(16)). For a fair comparison against p-IgGen(6), in which test sequences are selected on 95% concatenated CDR sequences across both chains, we created the “comparison_ds”. We clustered our data on 95% concatenated heavy chain CDR sequences and sampled 400 sequences different from training and validation, and present in both the test set of LICHEN and p-IgGen.

### Model architecture

LICHEN is a sequence-to-sequence model implemented according to the transformer from Vaswani et. al(17) in PyTorch(18). LICHEN consist of 6 encoder and decoder layers, all with 8 heads. The embedding size and the number of hidden dimensions were both set to 512, resulting in a total number of parameters of 25,284,632. Parameters were initialised using a Xavier normal distribution (Glorot initialization)(19).

The antibody sequences were tokenised using one-hot encoding of the amino acids as used in AbLang(15) and a start and stop token.

### Training protocol

LICHEN was trained using sinusoidal positional encoding for 4 epochs. The model was optimised using the Adam optimizer with default parameters except for the learning rate. We used a cosine scheduled learning rate with 5% warm-up steps and a maximum learning rate of ^10−4^, and a batch size of 64, a dropout of 0.1, and cross entropy loss. Training was performed on a single Quadro RTX 6000 GPU.

### Model inference

Light sequences were generated with a top-p parameter of 0.9 and a temperature parameter of 1.0. Minimal tuning of the two parameters showed that these values generate diverse and realistic light sequences.

LICHEN allows germline and CDR sequence information to be provided during inference. Users can restrict the model to generate a specific type (kappa or lambda), V-gene family (e.g. IGKV1), or V-gene (IGKV1-39). LICHEN is therefore seeded with either the first two (for type) or ten (for Vgene family and V-gene) residues. Initial residues are sampled based on observed frequency in germline sequences as stored in IMGT^2^. Users can also provide sequences of any length as a seed.

LICHEN can incorporate sequences of all three or a subset of the CDRs according to either the Kabat(20) or IMGT(21) definitions. As the architecture conditions the selection of the next residue on both the heavy sequence and previously generated light residues, LICHEN conditions on provided additional germline and CDR data. Correct CDR placement is established based on conserved residues and is checked by ANARCII(9). When CDR1 and CDR2 are provided without a light chain type, LICHEN is automatically seeded with the kappa or lambda type based on the closest alignment score of the CDRs against germline CDR residues using BLOSUM62(22). CDR1 is placed based on the conserved cysteine in IMGT position 23. Placing of conserved tryptophan in IMGT position 41 is forced after CDR1. CDR2 is placed based on the CDR1 or tryptophan 41, and the resulting sequence thus has a fixed FR2 length. CDR3 is placed according to the conserved cysteine in IMGT position 104 and subsequently a phenylalanine is forced after CDR3.

Generated light sequences for a specific heavy sequence can be automatically filtered by LICHEN. The filters implemented include: identification by ANARCII(9), human by Humatch(23), and most likely by AbLang2(24). Note that filtering on AbLang2 will result in germline diverse sequences. Moreover, filtering can be applied to select the most variable set of generated sequences and remove redundant sequences.

### Model evaluation

Light sequences generated by LICHEN should be valid, diverse, and a fit for the given heavy sequence.

The validity of generated sequences was checked by the ability of ANARCI(16) and ANARCII(9) to number the sequence and classify the sequence as a light sequence. Humanness of the sequence was checked using Hu-mAb(25) and Hu-match(23). The full VH and CDR sequence lengths were compared against the native paired light sequences. Combined CDR sequence length was compared against all native CDR lengths in the dataset. Structural diversity was analysed using canonical form assignment by SCALOP(26). ABodyBuilder2 (ABB2)(27) was used to structurally model the sequences.

The diversity of the generated sequences was evaluated by germline assignment and germline identity score as given by ANARCI. Sequence-based similarity of generated sequences to the training and validation data was determined by KAsearch(28). The structural diversity was evaluated on the structural_ds by canonical form assignment of CDR sequences using SCALOP, VH-VL orientation of ABB2 models by ABangle(29), and CDRH3 RMSD of ABB2 models. The latter was calculated by aligning the full VH sequence of the ABB2 antibody models with the 20 native or generated light sequences to their mean.

The fit of the generated light sequences for the heavy chain was evaluated based on the co-evolutionary relationship between the VH and VL arising from the affinity maturation process for the cognate antigen. This was checked based on the correlation between the number of mutations in the VH and VL, as well as the model preference for native pairing. For the former ANARCI germline identity scores were used. For the latter pairing probability scores were calculated on the conditioning_test. The conditional probability of the native light sequence given the native heavy sequences was normalised by the unconditional probability. This normalised probability was then compared to the normalised probability of the germline reverted light sequence given the native heavy sequence. Probability scores were calculated based on next token logits.

Model performance of LICHEN and p-IgGen were compared on the basis of ten generated light sequences for all heavy sequences in the comparison_ds. p-IgGen was therefore seeded with the heavy sequence.

### Experimental validation design

We also evaluated LICHEN experimentally by using it to predict sets of light chains for the therapeutics adalimumab(30) and pembrolizumab(31). Adalimumab is a fully human antibody (IGHV3-IGKV1) targeting TNF-α, a pro-inflamatory cytokine. Pembrolizumab is a humanised antibody (IGHV1IGKV3) targeting programmed death receptor 1 (PD-1). Three different use cases of LICHEN were evaluated: a case study with known CDRs (pairing_CDRs), a case study with the essential CDRs (pairing_eCDR), and a pairing case without CDR knowledge (pairing). The CDRL3 was assumed to be an essential CDR. For pembrolizumab the CDRL1 was also classified as essential due to its rare sequence length. For all cases, LICHEN was conditioned to make solely IGKV1, IGKV2, and IGKV3 light sequences for the native heavy sequences as these are most common.

Generated sequences were filtered on humanness by Humatch(23), ANARCI(16), and Hu-mAb(25). Sequences with known canonical forms according to SCALOP(26), and that could be modelled with ABB2(27), were selected. Developability was tested using TAP(32) and only sequences with five green flags were selected for pembrolizumab. For adalimumab four green flags and a minimum score for the structural Fv charge symmetry parameter SFvCSP equal to native adalimumab (−19.5) were allowed. For both therapeutic antibodies five light sequences were selected for the “pairing_CDRs” case, eight for the “pairing_eCDR” case, and ten for the “pairing” case (see Figure S1). For all three cases we selected diverse sequences.

The native light sequence was used as a positive control. Germline encoded V-gene sequences combined with Jgene IGKJ1*01 were used as baseline. IGKV1-39*01 and IGKV1D-33*01 were chosen as baseline for adalimumab IGKV3-20*01 and IGKV3-11*01 for pembrolizumab based on known binding preferences(33). The more rarely observed germline sequence IGKV7-3*01 was additionally used as a baseline. Baseline sequences were tested with and without native CDRs grafted for comparison to the CDR informed and standard pairing cases.

### Experimental validation

DNA coding for the amino acid sequence of each antibody was synthesised and cloned into the mammalian transient expression plasmid pETE V3 (property of Fusion Antibodies plc). Antibodies were expressed using a CHO-based transient expression system and the resulting antibodies were clarified by centrifugation and filtration. Antibodies were purified (using state-of-the-art AKTA chromatography equipment) from cell culture supernatants via affinity chromatography and analysed via size exclusion chromatography (SEC). The purity of the antibodies was determined to be >95%, as judged by reducing and denaturing Sodium Dodecyl Sulfate Polyacrylamide gels. SEC was run using a Superdex 200 Increase 10/300 GL column. Antibody concentration was determined by measuring absorbance at 280 nm and calculated using the standard extinction coefficient 205,500 *M*^*−*1^*cm*^*−*1^ (or 1.0 mg/ml = A280 of 1.37 [assuming a MW = 150,000 Da]) for an antibody.

A screening ELISA was prepared for all antibody variants created by LICHEN while conditioning on CDR data and controls that expressed and purified well. All antigens used in ELISA screening were purchased from Acro Biosciences. Purified TNF-α (Cat #TNA-H82E3, Lot #BV2043-241DF1-1R8) was used to test adalimumab and Human PD-1 (Cat #TNA-H82E3, Lot #BV2043-241DF1-1R8) was used to test pembrolizumab. A negative antigen (SARS-CoV-2 Spike RBD protein, Cat #SPD-C82E9, Lot #BV3541b-2043F1RD) was included to ensure binding specificity. Antigens were diluted to a concentration of 1 µg/mL in bicarbonate ELISA coating buffer and 30 µL (30 ng/well) was added to each well of a 384 well MaxiSorp plate (Nunclon). Plates were then incubated overnight at 4°C. Plates were washed 5 times with phosphate buffered saline (PBS) + 0.1% Tween using a Zoom HT plate washer (Berthold Technologies) and blocked with PBS containing 4% non-fat dry milk for 2 hours at room temperature, with shaking. All antibodies were diluted to a concentration of 1 µg/mL in PBS and 30 µL was added to each well (30 ng/well). Diluted antibodies were added to the plates with shaking for 2 hours at room temperature. Additional wells were coated with antigen and PBS was added in place of the adalimumab/pembrolizumab antibodies. Following sample incubation, plates were washed 5 times in PBS + 0.1% Tween using the Zoom plate washer before 1 hour incubation in Goat anti-human IgG – Fc specific HRP secondary antibody (1:70,000 dilution, Sigma A0170). Plates were washed a final time as before and 40 µL TMB II solution (Biopanda reagents) was added for 10 minutes at 37°C. The reaction was terminated by adding 20 µL of 1M Hydrochloric acid per well. Absorbance at 450 nm was then assessed using a Clariostar plate reader (BMG Labtech).

Variants that produced a positive signal (>0.2 nm at 450 nm) with the relevant antigen during ELISA analysis were selected for further binding analysis by Biolayer interferometry (BLI). A single-point analysis assay was performed by capturing the biotinylated TNF-α at 0.53 µg/mL and PD-1 at 0.28 µg/mL using SAX2 biosensors. The antigen-captured biosensors were submerged in wells containing each antibody, prepared at a single concentration of 33 nM (association stage), followed by a dissociation step in running buffer. To allow for double reference correction, antigen-captured sensors were dipped into wells containing only buffer. Blank sensors (no antigen present) were also dipped into wells containing each antibody. This referencing provided a means of compensating for any non-specific binding of the antibody to the sensor surface and also any baseline drift in the running buffer. Steps were performed at 25°C at a constant flow-rate of 1000 rpm. New sensors were used for each sample. All consumables used were those recommended by Sartorius. All samples were diluted in freshly prepared running buffer. Antigens were immobilized onto the surface of biosensors using the capture methods described previously. Antibody variants were passed over the surface to generate a binding response. Binding data for the Antibody:Antigen interactions were collected at 25°C on the biosensors.

## Results

LICHEN is a sequence-to-sequence model trained on human Fv sequences which generates antibody light sequences conditioned on the heavy chain. We evaluated generated light sequences based on validity, diversity, and fitness for the given heavy sequence. We experimentally validated LICHEN on a pairing use case and two pairing while maintaining antigen-binding use cases.

### LICHEN generates valid human light sequences

For the 20 generated light sequences for each of the 500 heavy sequences in the standard_test set we evaluated their antibody light chain likeliness. All generated sequences were identified by ANARCII as light sequences and could be numbered. Almost all sequences are human according to HumAb (99.92%) and Humatch (99.73%) and could be structurally modelled with ABB2 (99.80%) (see Table S1).

The VH and CDR sequence lengths show similar distributions to those observed in native light sequences (see Figure S2).

### Generated sequences are diverse

The diversity of sequences generated by LICHEN is in line with the variation observed in nature. According to germline assignment by ANARCI, generated sequences are closest to a diverse set of light chain V and J-genes and contain a varying number of mutations away from these germlines (see Figure S4). On average, generated sequences by LICHEN contain more mutations away from the germline V-gene compared to the native light sequences (see Figure 1). The canonical forms the structural conformation of the CDRs are equally diverse (see Figure S3).

**Figure 1.**
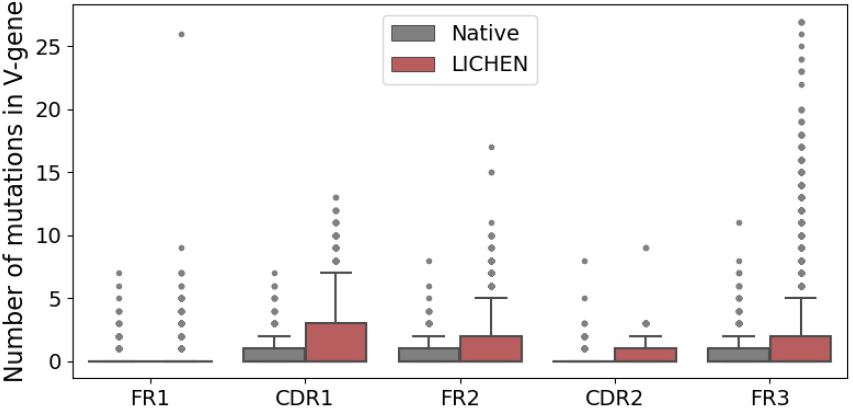
Number of mutations from the germline V-gene sequence per light sequence region. The native (grey) and LICHEN’s generated (red) light sequences of the standard_ds were compared against the closest germline V-gene according to germline annotations by ANARCI(16). Generated sequences by LICHEN contain in general more mutations away from the germline sequence.

Diversity is not only observed in light sequences generated for diverse heavy sequences, but also for a single heavy sequence (see Figure S5). We calculate the structural diversity of the light sequence CDRs using canonical form assignment, the Fv by the VH-VL orientation, and the unaltered CDRH3 sequence due to VL pairing by RMSD calculations. Again, the diversity is in line with the diversity of light sequences naturally pairing with the same heavy sequence (see Figure S5).

In general, LICHEN is more likely to generate light sequences of most frequent light chain types in the training data (see Figure S4). However, these sequences are not just germline sequences and LICHEN is able to generate novel full Fv, CDR and CDRL3 sequences compared to light sequences used for training and validation (see Figure S6).

### LICHEN has learned the co-evolutionary relationship between heavy and light sequences

It is known that during the antibody affinity maturation process sequences undergo somatic hypermutations in both chains. Optimised antibodies for the antigen are therefore expected to have a high number of mutations in both the heavy and light sequence. Figure 2 shows this co-evolutionary relationship between the VH and VL chain of the native sequences based on the average germline identity score of both the V and J-gene according to ANARCI. This relationship is not observed when randomly pairing the native heavy and light chains. Generated light sequences show a similarly strong correlation, indicating that LICHEN has captured this co-evolutionary relationship.

**Figure 2.**
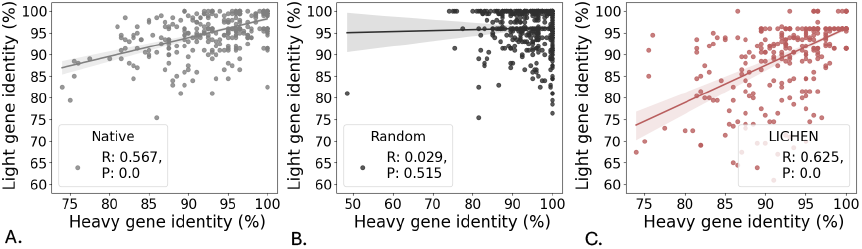
The average germline identity scores of the V-gene and J-gene derived from ANARCI(16) calculated for both chains of the native antibodies (A), random paired native antibodies (B), and antibodies with generated light sequences by LICHEN (C) on the standard_ds. A linear regression line, determined by SciPy(34), is shown as well as the corresponding Pearson correlation coefficient (R) and pvalue (P). LICHEN captured the co-evolutionary relationship between the heavy and light chain during affinity maturation.

We next tested if LICHEN also prefers pairing of the optimised memory heavy sequence with the optimised memory light sequence over pairing with the germline-reverted light sequence using the conditioning_ds. Figure 3 shows native pairings are preferred over germline-reverted pairings with a stronger preference observed when the light sequence contains at least ten mutations away from the germline.

**Figure 3.**
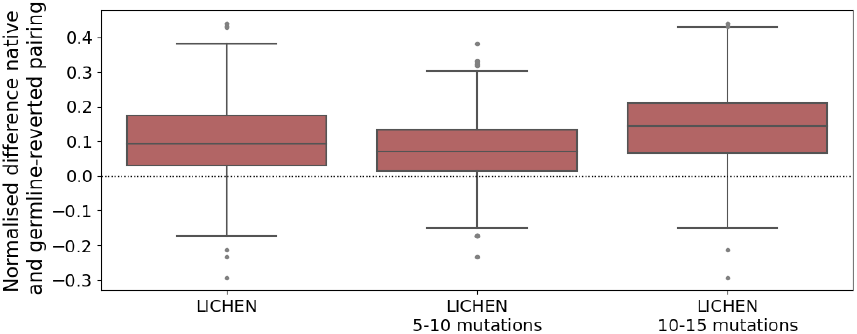
The normalised difference between the probability of native over germline reverted pairing. The conditional probability of generating the native light sequence and germline-reverted light sequence were normalised by the unconditional probability. The difference was normalised by the sum of the probabilities. Results are shown for the complete conditioning_ds and on subset containing either five to ten or ten to fifteenth mutations. Values above zero (dotted line) indicates the native pairing is preferred over the germline-reverted pairing. LICHEN in general prefers native pairing over germline-reverted pairings.

### LICHEN balances validity and diversity of generated light sequences

Diversity from the germline and fitness for the heavy sequence suggests LICHEN provides a valuable alternative to the standard strategy of taking a common germline light sequence to partner a heavy chain. We next compared LICHEN against an alternative machine learning model: p-IgGen(6). p-IgGen is a generative model for paired antibodies which can be seeded with initial sequential information; i.e. a light sequence can in principle be generated for a given heavy sequence. p-IgGen is not specifically trained for this task. Based on 10 generated light sequences for each of the 400 heavy sequences in the comparison_ds, p-IgGen is more likely to generate sequences that cannot be numbered and/or are not human according to *in silico* methods (see Table S2).

### LICHEN can be tailored to maintain key CDRs and select specific germlines

Existing generative models, such as p-IgGen, are unable to incorporate such additional information obtained prior to sequence design. LICHEN is designed to be able to incorporate binding information by conditioning on light chain CDRs. To accommodate for experimental choices on germline usage, e.g. when a specific Vgene is preferred based on prior knowledge of its characteristics or user expertise, LICHEN can be restricted to generation of a specific light chain type (kappa or lambda), V-gene family, or V-gene. We designed two *in vitro* case studies to demonstrate how LICHEN can be restricted to specific Vgene families and make use of CDR information for pairing.

### Sequences generated by LICHEN express well

For both the VH sequence of therapeutics adalimumab and pembrolizumab we generated VL sequences for three use cases: pairing_CDRs, pairing_eCDR, and pairing (see Methods). The generated sequences were evaluated *in silico*, after which a diverse set of sequences was selected for *in vitro* validation. Although adalimumab and pembrolizumab are optimised therapeutics, the light sequences generated by LICHEN and paired with the therapeutic VH exceeded the expression yield in 48% and 65% cases, respectively (Table S3). Only one antibody sequence generated by LICHEN for adalimumab, a IGKV2-24*01 light sequence, produced insufficient material. Reduced expression yields for pembrolizumab are observed when CDR information is incorporated for both LICHEN and baseline antibodies. The potential yield and monodispersity of all samples as well as the closest Vand J-gene of these sequences (according to ANARCI) are shown in Table S3. These results show that LICHEN can generate antibodies that are stable and express well.

### CDR tailored sequences generated by LICHEN maintain antigen-binding

Conditioning LICHEN on the light CDRs of adalimumab resulted in highly diverse sequences (see Figure S1) with strong expression yields while maintaining binding to TNF-α (see Figure 4B). Providing only CDRL3 (pairing_eCDR) created some weak binders. Similarly for pembrolizumab (Figure 4D), generated antibodies with strong expression yield maintain binding ability to PD1, and weak binders are observed among the paired_eCDR antibodies. Independent of expression yields, all germline sequences used as baselines maintain antigen-binding. The ELISA binding patterns are in line with BLI results, showing high on-rates and very slow dissociation for the high affinity binders (see Figure S7). These results indicate that when conditioning on binding information, LICHEN generates diverse light sequences with sensible expression yields that maintain target binding.

**Figure 4.**
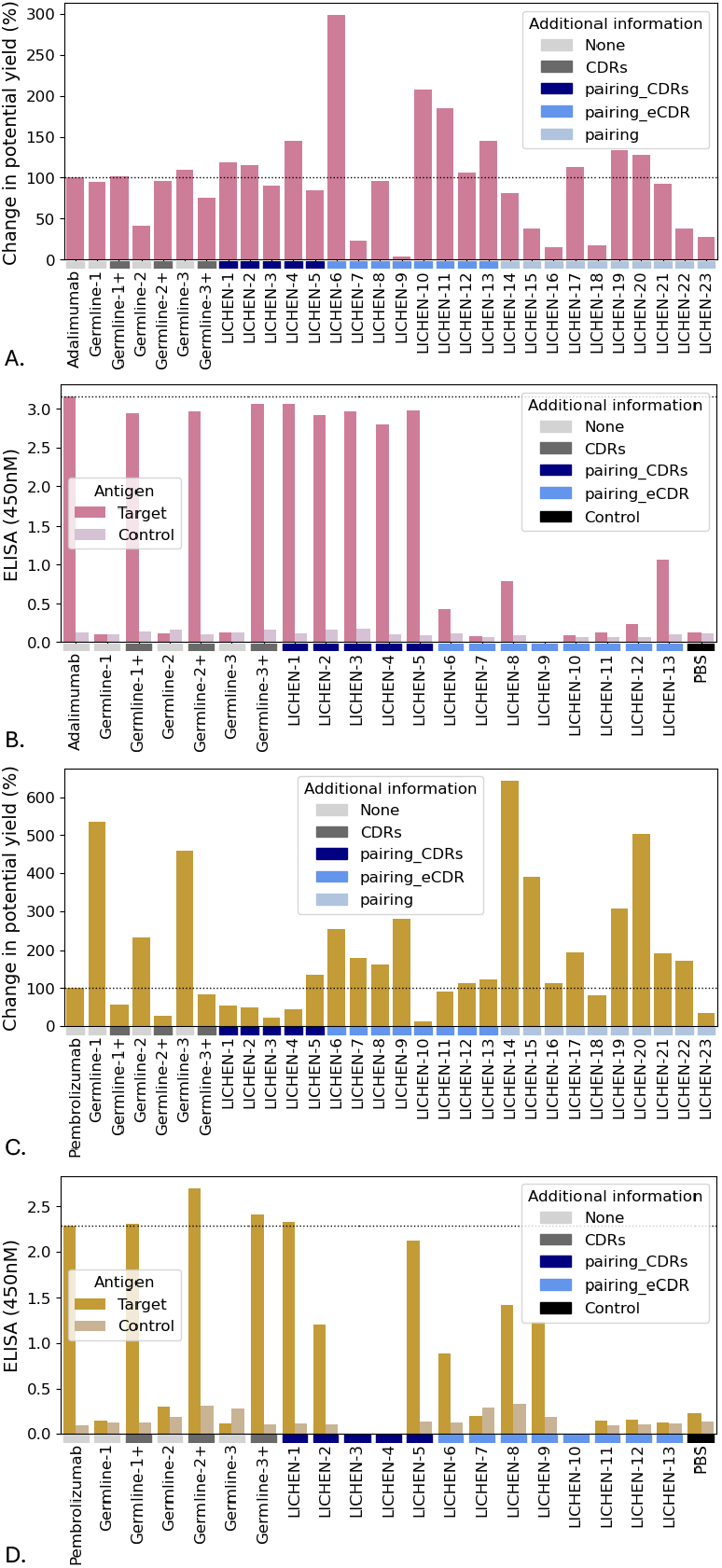
Percentage change in potential yield from therapeutic and binding data of generated light sequences by LICHEN for the VH of therapeutics adalimumab (A and B) and pembrolizumab (C and D). The therapeutics were used as positive controls. The germline V-gene sequences IGKV1-39*01, IGKV1D-33*01, and IGKV7-3*01 for adalimumab and IGKV3-20*01, IGKV3-11*01, and IGKV7-3*01 for pembrolizumab, labeld “Germline-1”, “Germline-2”, and “Germline-3” respectively, combined with IGKJ1*01 were used as baselines. All three light CDRs (Kabat definition) were grafted in the germlines labeled with the suffix “+” (dark grey). Three use cases where tested: pairing_CDRs (“LICHEN-1” to “LICHEN-5”, navy), pairing_eCDR (“LICHEN-6” to “LICHEN-13”, blue), and pairing (“LICHEN14” to “LICHEN-23”, light blue). For pairing_CDRs LICHEN was conditioned to all light therapeutic CDRs, for pairing_eCDR LICHEN was conditioned to essential light CDRs only (CDRL3 for adalimumab, CDRL1 and CDRL3 for pembrolizumab). LICHEN was restricted to make IGKV1, IGKV2, and IGKV3 light sequences and diverse sequences were tested. Binding was tested to the targets of adalimumab (TNF-α) and pembrolizumab (PD-1) (“Target”, pink and orange) and the unrelated non-human protein RBD was uses as a control (“Control”, light pink and light orange). Phosphate buffered saline (“PBS”, black) was used as secondary control. LICHEN generates well expressing antibodies which maintain binding ability to the target when additionally conditioned on the CDRs.

## Discussion

Generative machine learning models have shown success for antibody library design. However, methods are often explicitly designed for the initial exploration process and are purely computational. In practice, machine learning guidance and experimental design and validation go hand-in-hand.

To support the collaborative design of antibody therapeutics, we have developed LICHEN. LICHEN is a sequenceto-sequence model for light sequence generation. Light sequences are generated in a next-token fashion, conditioned on the heavy chain and preceding generated light chain residues.

LICHEN can also be conditioned by the user on CDRs and germline usage. This allows for generation of light sequences with the key CDRs for antigen binding. The three CDRs can be provided in any combination (e.g. only CDRL3) and according to both the IMGT(21) and Kabat(20) definitions. Limiting LICHEN to a specific light chain type (kappa or lambda), V-gene family, or V-gene allows for experimental choices.

This tailoring to experimental needs offers more practical applications compared to previous methods. LICHEN can be used in settings where only VH information or both VH and binding information is available. Furthermore, LICHEN allows two heavy sequences as input to find a common light sequence for a bispecific antibody. The automatic validation of generated sequences by LICHEN reduces the amount of required manual downstream analysis of the outputs.

Publicly available paired antibody data used for training LICHEN is biased towards specific V-genes, particularly to IGHV3, IGKV1, and IGKV3. Sequences generated by LICHEN show similar light chain biases. This preference is also observed in available therapeutic antibodies and was previously indicated as systematic bias in drug discovery pipelines(1).

Evaluating antibody generative models *in silico* is challenging, especially for heavy and light chain pairing. Heavy sequences have been shown to pair with diverse light sequences and vice versa, making the definition of an appropriate pairing unclear. Here, we constructed tests evaluating conventional light chain and pairing features. *In vitro* validations on two therapeutic VH chains confirmed that LICHEN generates diverse light sequences that are a fit in terms of expression and stability for these heavy chains.

During evaluation, we focused on finding an optimal balance between correctness and diversity of generated light sequences. Residues were selected using a top-p sampling approach allowing for generating diverse light sequences. The sequence and structural diversity generated by LICHEN for a given heavy sequence offers a broad exploration of the potential design space.

No increase in model performance was observed when increasing the number of parameters. This might be related to the relatively small amount of available paired antibody data. Extending the training data with the vast amount of unpaired sequence data did not improve performance. As more data became available after initially training LICHEN resulting in OAS storing more than 3M paired human sequences (June 2025) we retrained the model. Although the amount of data after filtering was increased (from 1.8M to 2.5M pairs), the data coverage is similar (see Figure S8) and no model improvement was observed.

To conclude, LICHEN generates human light sequences for a given heavy sequence. These light sequences are valid, diverse, and a fit for the heavy sequence. Additional tailoring towards key binding regions and restricting outputs to light chain types incorporates experimental needs. With experimental validations, we confirmed that the diverse light sequences created by LICHEN for two therapeutic VH chains form a stable and expressible antibody and, when conditioned on CDR information, maintain binding to the target. LICHEN thus offers a hybrid design strategy for diverse use cases at various stages in the antibody engineering workflow.

## Data availability

The filtered and cleaned data for training and testing have been deposited on Zenodo (https://doi.org/10.5281/zenodo.15917096).

## Code availability

LICHEN is available open source as both a python package (https://github.com/oxpig/LICHEN) and web application (https://opig.stats.ox.ac.uk/webapps/lichen).

## AUTHOR CONTRIBUTIONS

CMD and IE conceptualised the study. CMD, AGW, RJB, and CJM supervised the project. IE initialised model development. HLC continued model development, performed the research, and analysed the data. CJM prepared sequences for experimental validation. PD performed minipreps. GM performed the transient gene expression. MB and BC performed purification, SEC, and Quality Control. CB performed the ELISA and ST the BLI analysis. AGW developed the web tool. The manuscript was written by HLC and reviewed by RJB and CMD.

## ACKNOWLEDGEMENTS

The authors thank Benjamin H. Williams for his guidance on designing the web tool, and Lorna M. Stewart for helpful conversations on antibody design.

## Supplementary

**Figure S1.**
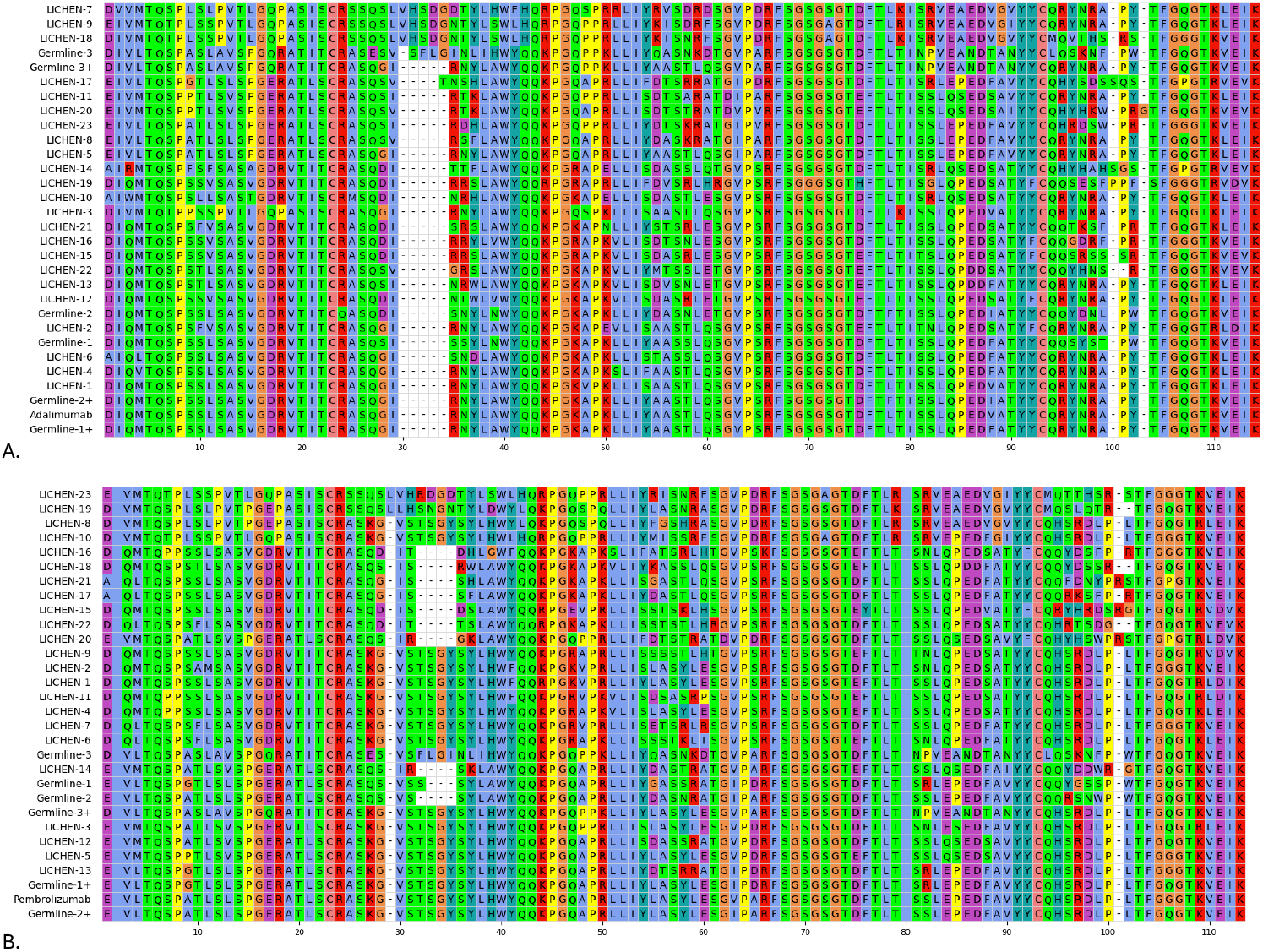
Generated light sequences by LICHEN were experimentally validated on the VH of therepautics adalimumab (A) and pembrolizumab (B). The therapeutics were used as positive control and germline V-gene sequence combined with IGKJ1*01 (Germline-1/3) as baseline. For adalimumab V-genes IGKV1-39*01, IGKV1D-33*01, and IGKV7-3*01 and for pembrolizumab IGKV3-20*01, IGKV3-11*01, and IGKV7-3*01 were used respectively. The “+” suffix indicates the therapeutic light CDRs (Kabat definition) were grafted into the germline sequences. Three use cases where tested: pairing_CDRs (LICHEN-1 to LICHEN-5), pairing_eCDR (LICHEN-6 to LICHEN-13), and pairing (LICHEN-14 to LICHEN-23). For pairing_CDRs LICHEN was conditioned to all light therapeutic CDRs, for pairing_eCDR LICHEN was conditioned to essential light CDRs only (CDRL3 for adalimumab, CDRL1 and CDRL3 for pembrolizumab). LICHEN was restricted to make only IGKV1, IGKV2, and IGKV3 light sequences. Generated sequences were *in silico* filtered based on ANARCI(16), Humatch(23), Hu-mAb(25), SCALOP(26), ABB2(27), and TAP(1, 32) after which 20 sequences were randomly chosen. Out of those, diverse sequences were selected for experimental validation.

**Table S1.**
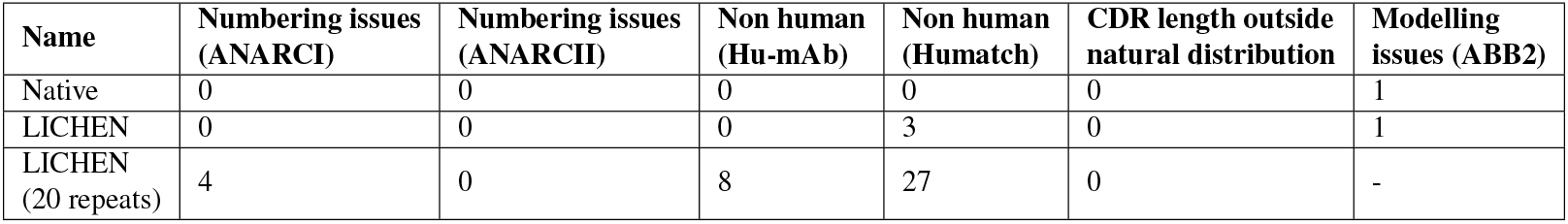
LICHEN generates valid (antibody-like) light sequences. For all entries in the standard_ds one (“LICHEN”) and 20 (“LICHEN (20 repeats)”) generated light sequences by LICHEN were validated based on the ability of ANARCI(16) and ANARCII(9) to number the sequence, humanness according to Hu-mAb(25) and Humatch(23), the number of CDRs outside a natural distribution and the ability of ABB2(27) to structurally model the sequence. Results are compared against the native light sequences.

**Table S2.**
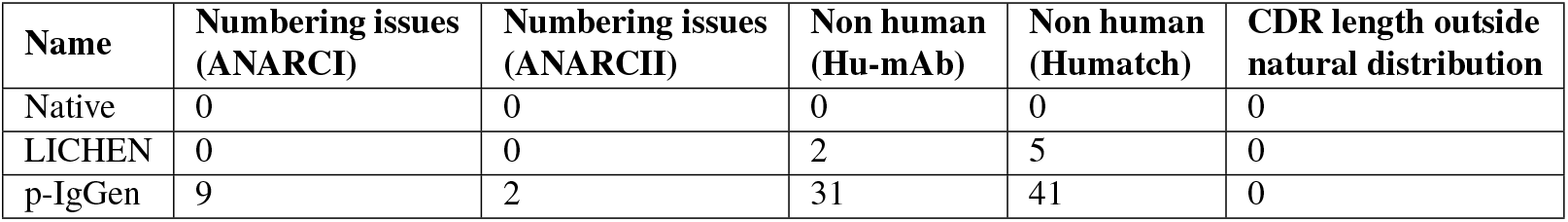
Validity of generated light sequences by LICHEN and p-IgGen(6) compared to the native light sequences in the comparison_ds. Sequences are validated based on the ability of ANARCI(16) and ANARCII(9) to number the sequence, humanness according to Hu-mAb(25) and Humatch(23), and the number of CDRs outside a natural distribution.

**Figure S2.**
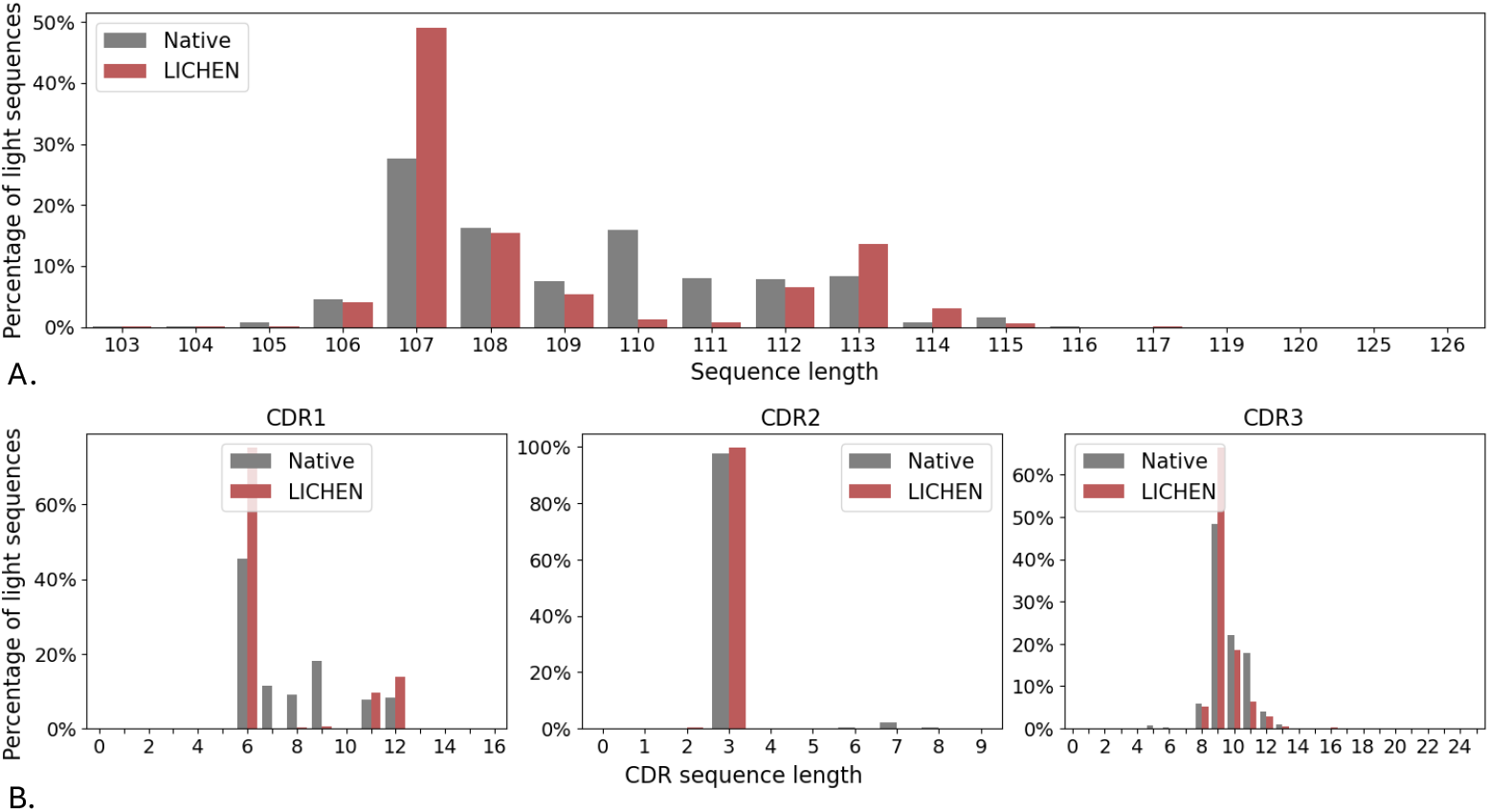
The full VL (A) and light CDR (B) sequence length of all the generated light sequences by LICHEN (red) compared to the native light sequences (grey) in the standard_ds. The percentage of light sequence with a certain length are shown.

**Figure S3.**
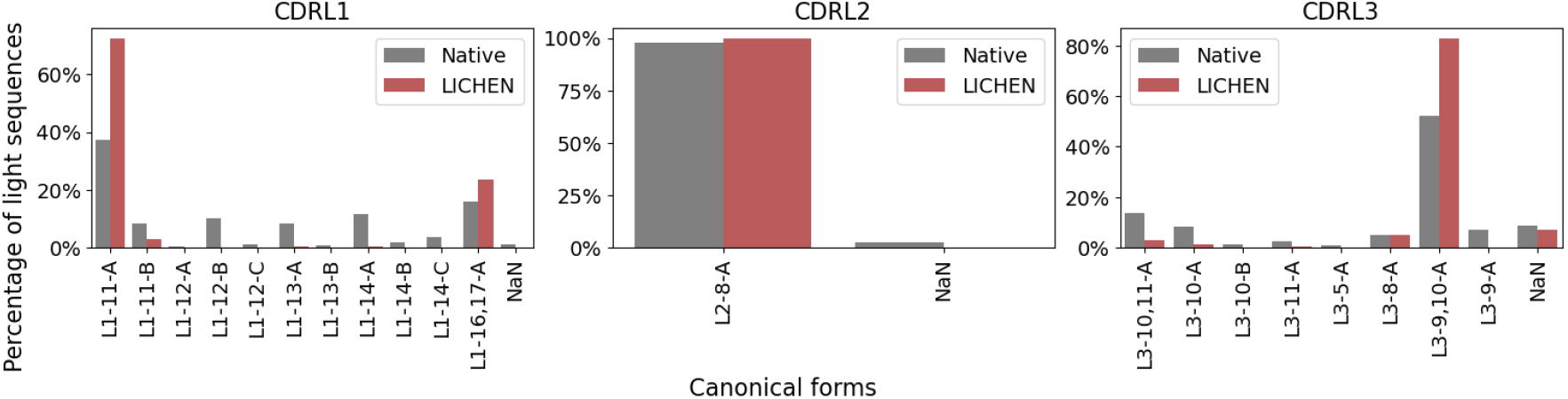
The canonical form assignment by SCALOP(26) on all generated light sequences by LICHEN (red) compared to the native light sequences (grey) in the standard_ds. The percentage of light sequence with a certain length are shown.

**Figure S4.**
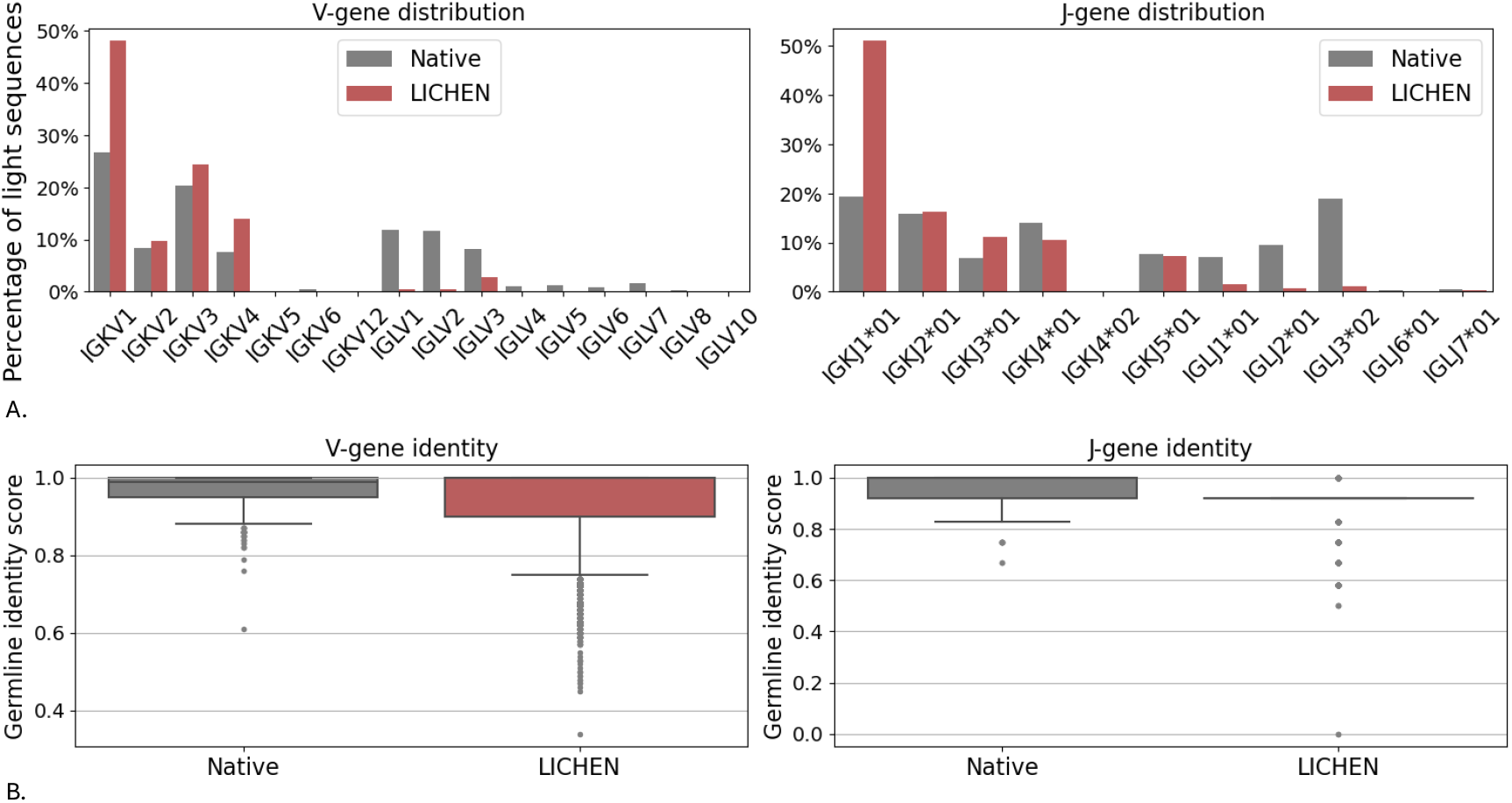
The germline V-gene family and J-gene diversity (A) and the corresponding germline identity score (B) of the generated light sequences by LICHEN (red) are compared against the native sequences (grey) in the standard_ds. The closest germline sequences and identity scores were determined by ANARCI(16).

**Figure S5.**
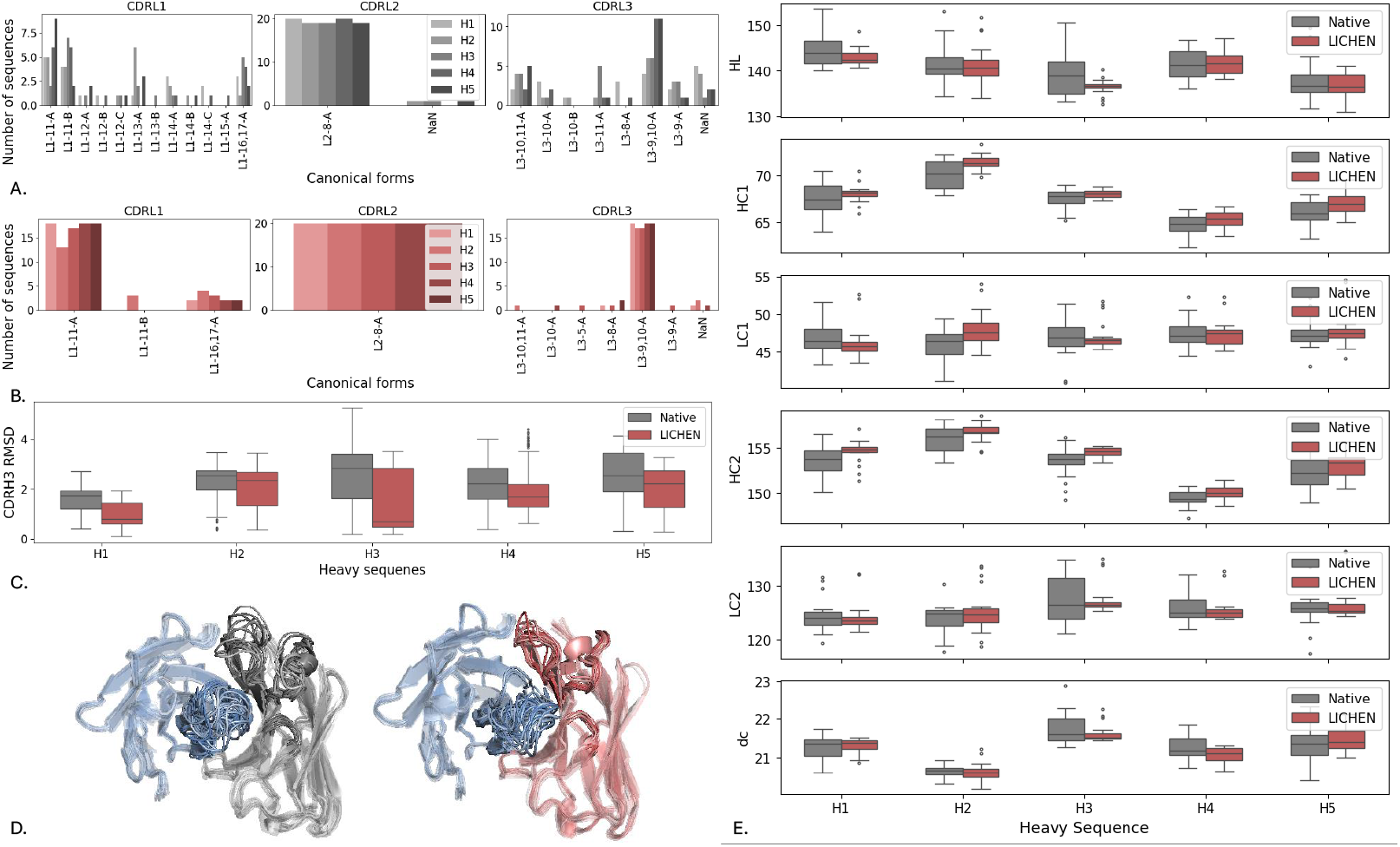
Structural diversity of twenty generated (red) and native (grey) light sequences pairing with five different heavy sequences of different V-genes: IGHV3 (H1), IGHV6 (H2), IGHV5 (H3), IGHV1 (H4), and IGHV4 (H5). (A and B) The canonical form assignment by SCALOP(26) (C) The RMSD of the unaltered CDRH3 sequences upon light sequence pairing. (D) Structural diversity of light chain CDRs (red) and CDRH3 (blue) of H4 are visualised with Pymol(35). (E) Variation in VH-VL orientation determined by ABangle(29).

**Figure S6.**
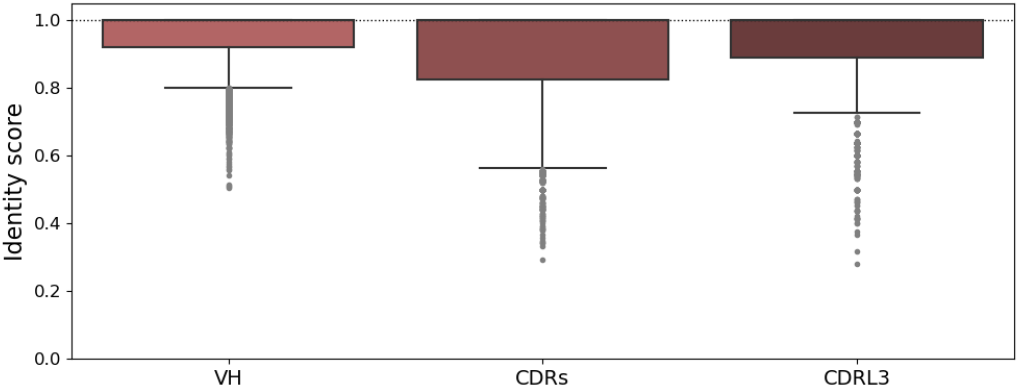
The identity score of the generated light sequence by LICHEN for the standard_ds to the training and validation set as determined by KAsearch(28). Identity scores are shown for the full VH, light chain CDRs, and the CDRL3.

**Table S3.** The V-gene and J-gene (determined by ANARCI(16)), potential yield, and monodispersity of the experimental validated generated sequences by LICHEN and the control sequences. Monodispersity is based on a SEC analysis.

| Adalimumab |  |  |  |  | Pembrolizumab |  |  |  |
| --- | --- | --- | --- | --- | --- | --- | --- | --- |
| Name | V-gene | J-gene | Potential yield (mg/L) | Monodispersity (%) | V-gene germline | J-gene | Potential yield (mg/L) | Monodispersity (%) |
| Therapeutic | IGKV1-27*01 | IGKJ1*01 | 53 | >95 | IGKV3-11*01 | IGKJ4*01 | 32 | >95 |
| Germline-1 | IGKV1-39*01 | IGKJ1*01 | 50 | >90 | IGKV3-20*01 | IGKJ1*01 | 171 | >95 |
| Germline-1+ | IGKV1-39*01 | IGKJ1*01 | 54 | >95 | IGKV3-20*01 | IGKJ1*01 | 18 | >95 |
| Germline-2 | IGKV1D-33*01 | IGKJ1*01 | 22 | >90 | IGKV3-11*01 | IGKJ1*01 | 75 | >95 |
| Germline-2+ | IGKV1D-33*01 | IGKJ1*01 | 51 | >95 | IGKV3-11*01 | IGKJ1*01 | 9 | >95 |
| Germline-3 | IGKV7-3*01 | IGKJ1*01 | 58 | >90 | IGKV7-3*01 | IGKJ1*01 | 147 | >95 |
| Germline-3+ | IGKV7-3*01 | IGKJ1*01 | 40 | >95 | IGKV7-3*01 | IGKJ1*01 | 27 | >95 |
| LICHEN-1 | IGKV1-27*01 | IGKJ2*01 | 63 | >95 | IGKV1-27*01 | IGKJ5*01 | 17 | >60 |
| LICHEN-2 | IGKV1-27*01 | IGKJ5*01 | 61 | >95 | IGKV1-17*03 | IGKJ4*01 | 16 | >40 |
| LICHEN-3 | IGKV1-27*01 | IGKJ2*01 | 48 | >95 | IGKV3-11*01 | IGKJ5*01 | 7 | - |
| LICHEN-4 | IGKV1-16*01 | IGKJ2*01 | 77 | >95 | IGKV1-39*01 | IGKJ2*01 | 14 | - |
| LICHEN-5 | IGKV3-11*01 | IGKJ2*01 | 45 | >95 | IGKV3-15*01 | IGKJ4*01 | 43 | >95 |
| LICHEN-6 | IGKV1-13*02 | IGKJ2*01 | 158 | >95 | IGKV1-9*01 | IGKJ2*01 | 81 | >95 |
| LICHEN-7 | IGKV2-30*01 | IGKJ2*01 | 12 | >95 | IGKV1-9*01 | IGKJ4*01 | 57 | >95 |
| LICHEN-8 | IGKV3-11*01 | IGKJ2*01 | 51 | >95 | IGKV2-28*01 | IGKJ5*01 | 52 | >90 |
| LICHEN-9 | IGKV2-24*01 | IGKJ2*01 | 2 | - | IGKV1-39*01 | IGKJ3*01 | 90 | >85 |
| LICHEN-10 | IGKV1D-8*02 | IGKJ2*01 | 110 | >95 | IGKV2-24*01 | IGKJ4*01 | 4 | - |
| LICHEN-11 | IGKV3-15*01 | IGKJ2*01 | 98 | >95 | IGKV1-16*01 | IGKJ5*01 | 29 | >65 |
| LICHEN-12 | IGKV1-5*01 | IGKJ2*01 | 56 | >95 | IGKV3-15*01 | IGKJ4*01 | 36 | >85 |
| LICHEN-13 | IGKV1-5*01 | IGKJ2*01 | 77 | >95 | IGKV3-20*01 | IGKJ4*01 | 39 | >90 |
| LICHEN-14 | IGKV1-8*01 | IGKJ3*01 | 43 | >95 | IGKV3-15*01 | IGKJ1*01 | 206 | >90 |
| LICHEN-15 | IGKV1-12*01 | IGKJ1*01 | 20 | >90 | IGKV1-27*01 | IGKJ3*01 | 125 | >95 |
| LICHEN-16 | IGKV1-33*01 | IGKJ4*01 | 8 | >95 | IGKV1-16*01 | IGKJ4*01 | 36 | >80 |
| LICHEN-17 | IGKV3-20*01 | IGKJ3*01 | 60 | >90 | IGKV1-13*02 | IGKJ1*01 | 62 | >90 |
| LICHEN-18 | IGKV2-24*01 | IGKJ4*01 | 9 | >90 | IGKV1-5*03 | IGKJ1*01 | 26 | >55 |
| LICHEN-19 | IGKV1-12*01 | IGKJ3*01 | 71 | >95 | IGKV2-28*01 | IGKJ1*01 | 98 | >85 |
| LICHEN-20 | IGKV3-15*01 | IGKJ1*01 | 68 | >95 | IGKV3-15*01 | IGKJ3*01 | 161 | >95 |
| LICHEN-21 | IGKV1-12*01 | IGKJ2*01 | 49 | >90 | IGKV1-13*02 | IGKJ3*01 | 61 | >95 |
| LICHEN-22 | IGKV1-5*01 | IGKJ1*01 | 20 | >85 | IGKV1-9*01 | IGKJ1*01 | 55 | >95 |
| LICHEN-23 | IGKV3-11*01 | IGKJ4*01 | 15 | >90 | IGKV2-24*01 | IGKJ4*01 | 11 | >95 |

**Figure S7.**
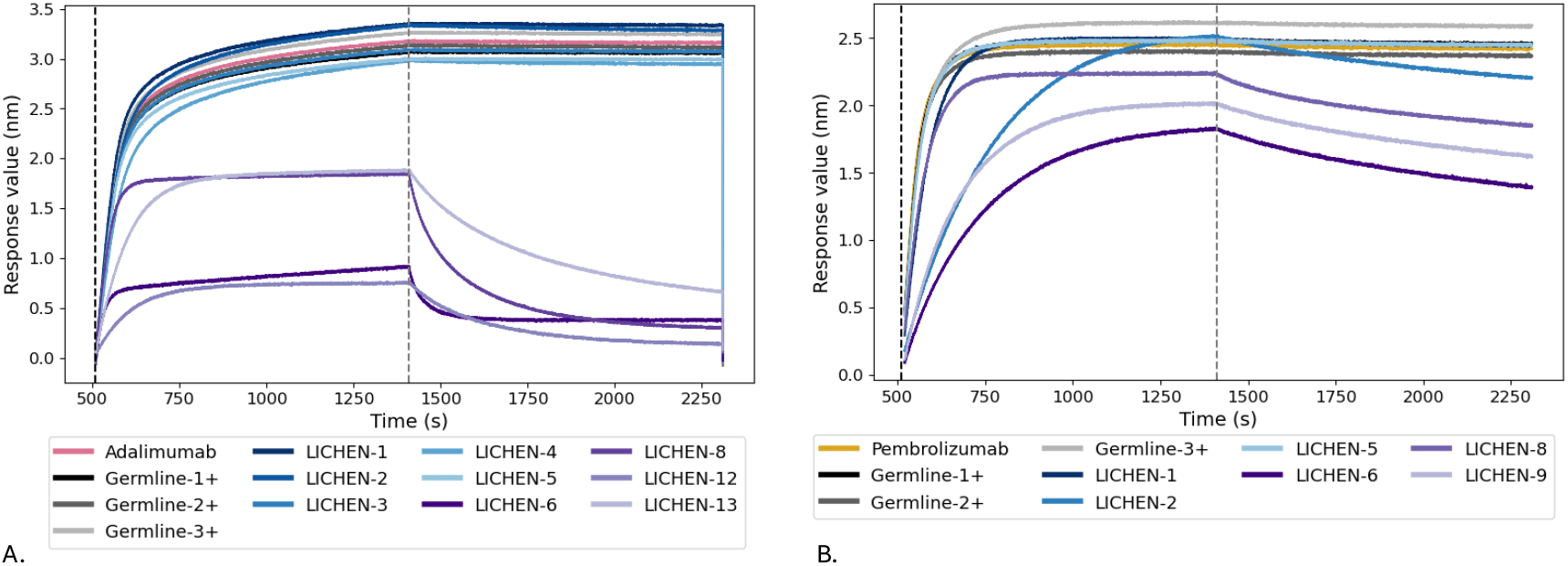
Bio-layer interferometry analysis of the therapeutic antibodies adalimumab (A, pink) and pembrolizumab (B, orange), and the control and generated light sequences pairing with the therapeutic VH sequence showing effective target binding in ELISA. Germline light sequences with grafted CDRs are shown in a grey palette and light sequences generated by LICHEN in a blue palette (all CDRs; pairing_CDRs) and in a purple palette (essential CDRs; pairing_eCDR). The association phase is shown between the dotted black (510 s) and the dotted grey line (1410 s). The dissociation phase is shown after the dotted grey line.

**Figure S8.**
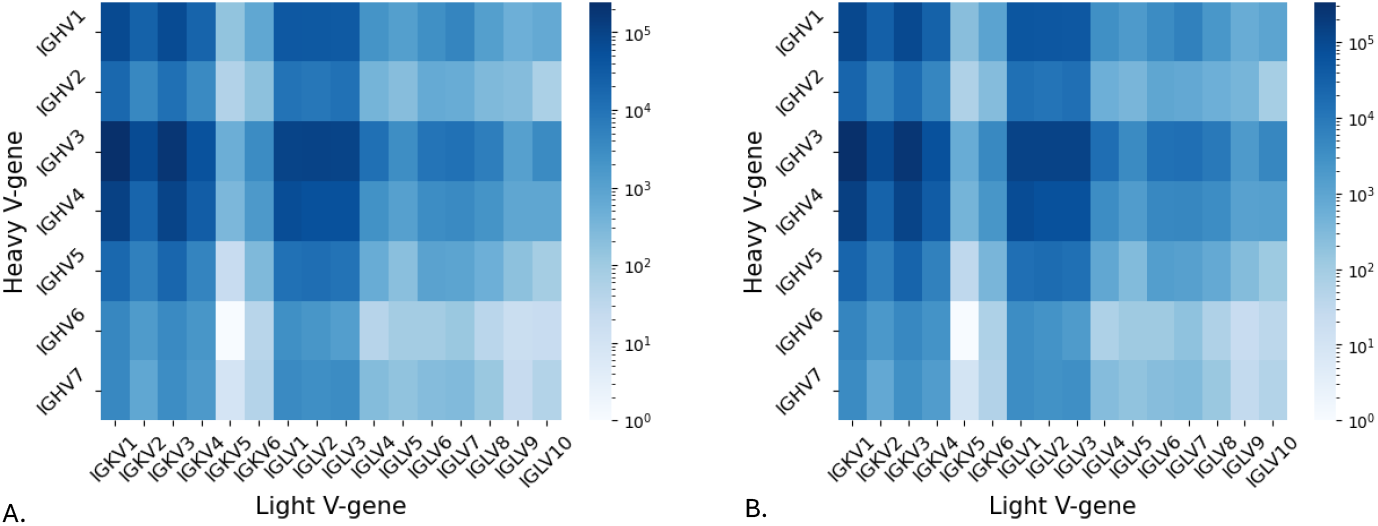
The distribution of V-gene of the heavy and light paired sequences stored in OAS(11) after filtering in November 2023 (A) and June 2025 (B). Similar gene distributions are observed for the 1.8M paired sequences as for the 2.5M paired sequences.

## Footnotes

1 https://opig.stats.ox.ac.uk/webapps/oas, assessed on 5 June 2025

2 https://www.imgt.org/genedb/, assessed in January 2025

